# Humans recognize affective cues in primate vocalizations: Acoustic and phylogenetic perspectives

**DOI:** 10.1101/2022.01.26.477864

**Authors:** C. Debracque, Z. Clay, D. Grandjean, T. Gruber

## Abstract

Humans are adept in extracting affective information from the vocalisations of not only humans but also other animals. Current research has mainly focused on phylogenetic proximity to explain such cross-species emotion recognition abilities. However, because research protocols are inconsistent across studies, it remains unclear whether human recognition of vocal affective cues of other species is due to cross-taxa similarities between acoustic parameters, the phylogenetic distances between species, or a combination of both. To address this, we first analysed acoustic variation in 96 affective vocalizations, including agonistic and affiliative contexts, of humans and three other primate species – rhesus macaques, chimpanzees and bonobos – the latter two being equally phylogenetically distant from humans. Using Mahalanobis distances, we found that chimpanzee vocalizations were acoustically closer to those of humans than to those of bonobos, confirming a potential derived vocal evolution in the bonobo lineage. Second, we investigated whether 68 human participants recognized the affective basis of vocalisations through tasks by asking them to categorize (‘A vs B’) or discriminate (‘A vs non-A’) vocalisations based on their affective content. Results showed that participants could reliably categorize and discriminate most of the affective vocal cues expressed by other primates, except threat calls by bonobos and macaques. Overall, participants showed greatest accuracy in detecting chimpanzee vocalizations; but not bonobo vocalizations, which provides support for both the phylogenetic proximity and acoustic similarity hypotheses. Our results highlight for the first time the importance of both phylogenetic and acoustic parameter level explanations in cross-species affective perception, drawing a more complex picture to explain our natural understanding of animal signals.

## Introduction

Vocal communication of affect is crucial for the emotional and attentional regulation of human social interactions (Grandjean et al., 2005; Sander et al., 2005; Schore & Schore, 2008). For instance, the modulation of prosodic features in human speech such as intonation or amplitude can convey subtle affective information to receivers (Grandjean, Bänziger, & Scherer, 2006; Scherer, 2003). Humans consistently recognize and evaluate the affective cues of others’ vocal signals in tasks with varying levels of complexity, with emotion categorization i.e. unbiased choice (A versus B) seemingly more cognitively complex than discrimination i.e. biased choice (A versus non-A) (Dricu et al., 2017; Gruber et al. 2020). In both emotion categorization and discrimination tasks, research shows that listeners can subjectively attribute the speaker’s reported affective state (i.e. angry, fearful or happy) as well as any potentially referential content (Brunswick, 1956; Grandjean et al., 2006). By no means uniquely human, these affective identification mechanisms facilitate adaptive behaviour in animals such as to approach or avoid the stimulus (Frijda, 1987, 2016; Gross, 1998; Nesse, 1990). Hence, current mechanisms underlying human and other animal vocalizations seem to result from similar adaptive pressures. For instance, research has shown the critical role of acoustic roughness in both human and great ape fear screams to rapidly appraise danger (Arnal et al., 2015; Kret et al., 2020). Despite the adaptive value and importance of auditory affective processing to our own species, its evolutionary origins remain poorly understood.

As noted, the adaptive behaviours underpinning communication of affect are often shared amongst animals. Over a century ago, Darwin (1872) hypothesized an evolutionary continuity between human and other animals for the vocal expression of affective signals. Morton (1977, 1982) subsequently proposed a model of motivational structural rules to characterize the relationship between the acoustic structure of mammal and bird vocalizations and their presumed affective contents. The systematic modulation of call acoustic structure and the caller’s underlying affective state appear to provide reliable cues that allow listeners to evaluate aspects of the eliciting stimulus, such as the level of threat or danger (Anderson & Adolphs, 2014; Filippi et al., 2017). Comparative research has confirmed that conspecifics are sensitive to such cues, with playback studies showing that both chimpanzees and rhesus macaques discriminate between agonistic screams produced by victims facing varying degrees of threat (Slocombe, Townsend, & Zuberbühler, 2009; Gouzoules, 1984), while meerkats extrapolate the degree of urgency required from the acoustic structure of conspecific alarm calls (Manser, 2001). This evidence suggests an evolutionary continuity in the vocal processing ability of both humans and non-human primates to accurately identify affective cues in conspecific vocalizations (Gruber & Grandjean, 2017).

Interestingly, this evolutionary continuity is also suggested by a second line of research, which shows that human participants generally perform above chance asked to identify primate signals. Despite a limited number of currently available studies (eight, to our knowledge - Belin, Fecteau, et al., 2008; Ferry et al., 2013; Filippi et al., 2017; Fritz et al., 2018; Kamiloğlu et al., 2020; Kelly et al., 2017; Linnankoski et al., 1994; Scheumann et al., 2014, 2017), existing findings on human perception of arousal and valence in non-human primate calls are promising. Indeed, research has shown that humans can discriminate the valence of chimpanzee vocalizations, including agonistic screams (negative valence) and food-associated calls (positive valence) (Fritz et al., 2018; Kamiloğlu et al., 2020); by comparison however, behavioural discrimination for rhesus macaque calls given in the same contexts is poor (Fritz et al., 2018; Belin et al., 2008). Functional Magnetic Resonance Imaging (fMRI) measures taken by Fritz and collaborators also showed that neural activations were more similar when attending to chimpanzee and human vocalizations than macaque calls. In contrast, Linnankoski and colleagues (1994) found that both human adults and infants could categorize affective macaque vocalizations in a larger range of contexts (angry, fearful, satisfied, scolding and submissive). Methodological differences might explain the differences in previous findings concerning macaque calls: it may be easier for human adults and infants to label affective contents of non-human primate vocalizations in a forced choice paradigm (categorization or discrimination tasks) in which the number of possibilities is limited rather than to rate the valence or arousal using Likert scales. For instance, research with human affective stimuli using forced choice paradigms demonstrated the positive relationship between cognitive complexity and the number of available categories to choose from (Dricu et al., 2017; Gruber et al. 2020). Thus, forced choice paradigms with limited options to choose from may lead to elevated performance with macaque calls (Linnankoski et al., 1994) compared to paradigms with Likert rating scales (Belin et al., 2008; Fritz et al., 2018).

In addition to the mixed findings concerning human sensitivity to valence in non-human primate vocalisations, evidence that humans can accurately judge vocal arousal in other species is also mixed. Recent findings highlight the ability of humans to reliably identify arousal in barbary macaque vocalizations expressed in negative contexts (Filippi et al., 2017) and arousal ratings of chimpanzee vocalizations seem to be fairly accurate across positive and negative valences (Kamiloğlu et al., 2020). Yet, Kelly and collaborators (2017) also showed that human participants over-estimated the distress content of bonobo infant calls compared to those of human or chimpanzee ones, suggesting a relatively poor capacity of humans to identify arousal in bonobo vocalizations. Overall, humans appear to perform relatively well with chimpanzee calls (Kamiloğlu et al., 2020), but less well with bonobo or macaque calls. However, it remains unclear why this is the case. In addition, it is also relevant to examine why a particular primate species, human especially, may be able to recognize affective vocalizations expressed by another primate species.

Several factors might explain our abilities to recognize some species’ affective vocalizations more reliably than others. Previous studies comparing human responses to closely and distantly related species, have highlighted the importance of phylogenetic proximity in human recognition of affect (e.g. Belin et al. 2008, Fritz et al. 2018), arguing that we are more sensitive to emotional content of vocalisations in closely related species. An important test of this hypothesis is to examine responses to vocalisations of two species that are equally closely related to humans. Only one study has attempted this to date by comparing human responses to chimpanzee and bonobo vocalisations, humans closest living relatives (Gruber & Clay, 2016). Focusing on distress calls, Kelly et al (2017) found that humans were less accurate at rating distress intensity in bonobo calls compared to chimpanzee calls, but whether this pattern generalizes beyond distress calls is currently unknown.

In addition to phylogenetic proximity, another important factor determining human accuracy at detecting the emotional content of other species vocalisations may be similarity in the acoustic parameters of vocalisations between humans and the test species. Previous studies have revealed cross-taxa similarities in the acoustic conveyance of affect (Ross, Owren, & Zimmermann, 2009; Scheumann et al., 2014). In particular, previous research has linked the human ability to recognize affective cues from vocalizations of other species to specific modulations of the fundamental frequency (F0), the mean pitch or the energy of the affective calls expressed by non-human primates (Briefer, 2012, Filippi et al., 2017; Linnankoski et al., 1994; Scheumann et al., 2014). Concurrently, acoustic similarity is also influenced by the call’s emotional valence (Belin, Fecteau, et al., 2008). Despite being as equally related to us as chimpanzees, the vocal repertoire of bonobos shows some notable acoustic differences, including elevated pitch (Tuttle, 1993) potentially due to shorter vocal tracts (Grauwunder et al 2018). Hence, it seems reasonable to hypothesize that acoustic differences in bonobo calls may lead to lower performance in a human recognition task.

Overall, it thus remains unclear whether the human ability to recognize affective vocal cues from other species is mainly due to (1) cross-taxa similarities in acoustic parameters, (2) the phylogenetic distances between species, or (3) both, considering that closely phylogenetically-related species may be likely to share acoustic parameters. To address these outstanding issues, we designed a forced-choice paradigm, where participants had to perform two tasks: categorization (A versus B, cognitively demanding) and discrimination (A versus non-A; less cognitively demanding). In both tasks, participants had to judge the affective nature of vocalisations produced in three affective contexts (threat, distress and affiliation) by humans and three other primate species that vary in phylogenetic distance to humans (equally close to humans: chimpanzee, bonobo; more distant: rhesus macaque). For each of the two tasks we measured whether participants were significantly above chance, and whether accuracy of performance could be predicted by species, affect or their interaction. To disentangle whether human cross-species emotion recognition performance was best explained by phylogenetic distance or acoustic similarity, we first established the acoustic similarity of chimpanzee, bonobo and macaque vocalisations to human vocalisations. We calculated Mahalanobis distances to compare the acoustic distances between vocalizations of various affective contexts from these species. We expected that if phylogenetic distance was the main determinant of performance, recognition of affective cues in human vocalisation should be greater than those of chimpanzees and bonobos, which should be equally better than those of rhesus monkey vocalizations (Humans>Chimpanzees=bonobos>macaques). By contrast, if acoustic similarity was the main determinant of performance, participants should perform best with the calls of species most acoustically similar to those of humans. If we found a significant interaction between species and affect on Mahalanobis distance of calls to the human centroid, then recognition performance would need to be compared to acoustic similarity between species at the level of each affect. Moreover, both phylogenetic proximity and acoustic distance may both play a role in explaining human cross species emotional recognition. We may expect amongst equally related species, more accurate performance with the species most similar acoustic structures to humans (if chimpanzees are shown to be more acoustically similar to humans than bonobos overall, or for certain affects, we might expect better recognition accuracy for chimpanzees than bonobos: Humans > Chimpanzees > Bonobos > Macaques). Finally, because of the previous literature (Dricu et al. 2017; Gruber et al. 2020), we also expected participants to perform more accurately on discrimination rather than categorisation tasks.

## Materials and methods

### Participants

Sixty-eight healthy adult volunteers from the Geneva area (29 males; mean age 23.54 years, SD = 5.09, age range 20 – 37 years) took part in the experiment. The participants reported normal hearing abilities and normal or corrected-to-normal vision. No participant presented a neurological or psychiatric history, or a hearing impairment. All participants gave informed and written consent for their participation in accordance with the ethical and data security guidelines of the University of Geneva. The study was approved by the Ethics Cantonal Commission for Research of the Canton of Geneva, Switzerland (CCER).

### Vocal stimuli

For our stimuli, we compiled a set of ninety-six vocalizations balanced across four primate species (human, chimpanzee, bonobo, rhesus macaque) and three affective contexts (threat, distress and affiliation). For human stimuli, non-linguistic vocal stimuli from two male and two female actors denoted as expressing a happy, angry or fearful affect were obtained from the Montreal Affective Voices Audio Collection (Belin, Fillion-Bilodeau, et al., 2008). For chimpanzee, bonobo and rhesus macaque stimuli, vocalizations taken from existing author databases were compiled from corresponding contexts: affiliation - food-associated grunts, threat - aggressor barks in agonistic contexts, and distress calls - victims in social conflicts. For each species, 24 stimuli taken from 6-8 different individuals were selected containing single calls or two call sequences of a single individual. All vocal stimuli were standardized to 750 milliseconds using PRAAT (www.praat.org) but were not normalized for energy to preserve the naturality of the sounds (Ferdenzi et al., 2013).

### Experimental procedure

Seated in front of a computer, participants listened to the vocalizations played binaurally using Seinnheiser headphones at 70 dB SPL. Each of the 96 stimuli was repeated nine times across six separate counterbalanced blocks leading to 864 trials following a randomization process. The overall experiment followed a within-subjects design with various layers (Figure 1). Testing blocks were task-specific, with participants either performing a categorization task (A versus B) or a discrimination task (A versus non-A). Participants completed three categorization blocks and three discrimination blocks, resulting in six blocks in total. Each block was made of 12 mini-blocks, each separated by a break of 10 seconds. Mini-blocks comprised one unique mini-block per species (human, chimpanzee, bonobo and rhesus macaque), each mini-block repeated 3 times. Within each mini-block were 12 trials, containing four vocalisations from all three contexts (affiliative/happy; threatening/anger; distress/fear) produced by a single species. The blocks, mini-blocks and stimuli were pseudo-randomly assigned for each participant to avoid more than two consecutive blocks, mini-blocks and stimuli from the same category.

**Figure 1:**
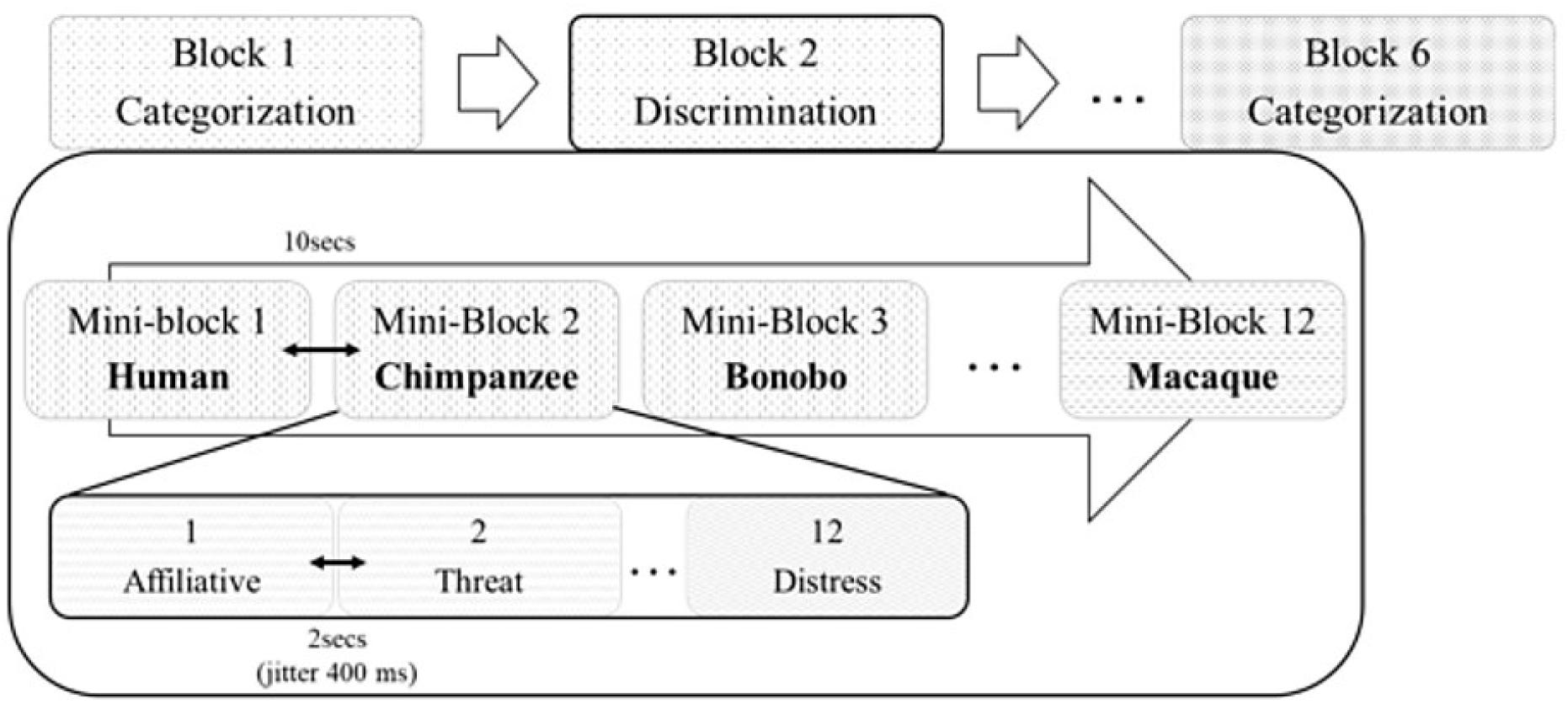
Structure of the experiment, with each of the six blocks made of 12 mini-blocks, which in turn comprised 12 individual trials.

At the beginning of each block, participants were instructed to identify the affective content of the vocalizations using a keyboard. For instance, the instructions for the categorization task could be “Affiliative – press M or Threatening – press Z or Distress – press space bar”. Similarly, the instructions for discrimination could be “Affiliative – press Z or other affect – press M”. The pressed keys were randomly assigned across blocks and participants. The participants pressed the key during 2-second intervals (jittering of 400 ms) between each stimulus. If the participant did not respond during this interval, the next stimulus followed automatically.

### Statistical analysis

#### Acoustic analyses

To quantify the impact of acoustic distance in human affect recognition of primate vocalizations, we automatically extracted 88 acoustic parameters from all stimuli vocalizations using the extended Geneva Acoustic parameters set, which is defined as the optimal acoustic indicators related to human voice analysis (GeMAPS; Eyben et al., 2016). This set of acoustical parameters was selected based on i) their potential to index affective physiological changes in voice production, ii) their proven value in former studies as well as their automatic extractability, and iii) their theoretical significance. This set of acoustic parameters includes related frequency parameters (e.g. pitch, jitter, formants), energy parameters (e.g. loudness, shimmer), and spectral parameters (e.g. alpha ratio, Hammarberg index, spectral slopes).

To assess the acoustic distance between vocalizations of all species, we then ran a Discriminant Analysis (DA) using SPSS 26.0.0.0 based upon the 88 acoustical parameters in order to discriminate our stimuli based on the four different species (human, chimpanzee, bonobo, and rhesus macaque). Excluding the acoustical variables with the highest correlations (>.90) to avoid redundancy of acoustic parameters, we retained 16 acoustic parameters related to frequency, energy, and spectral parameters that could discriminate species (see Supplementary material Table S1).

Using these 16 acoustic features, we subsequently computed Mahalanobis distances of the 96 experimental stimuli. A Mahalanobis distance is obtained from a generalized pattern analysis computing the distance of each vocalization from the centroids of the different species vocalizations (Mahalanobis, 1936). This analysis allowed us to obtain an acoustical distance matrix used to test how these acoustical distances were differentially related to the different species. To test this, we performed Generalized Linear Mixed Models (GLMMs) fitted by Restricted Maximum Likelihood (REML) on R.studio (Team, 2020) using the package Lme4 (Bates et al., 2015) to test whether the following three fixed factors could predict the Mahalanobis distances: Species (the species which produced the vocalization), Distance-Species (the species centroid used to compute the distance for the same species or for the other species, e.g. the human centroid used to quantify the distance of chimpanzee vocalization from humans), and Affect (affiliative, threat, and distress). We also examined the interaction between these three factors. The identity of the vocalizer was included as a random factor.

To test the effects of phylogenetic distance, we performed contrasts of interest on the factor of Species (i.e. human < chimpanzee=bonobo < macaque) taking into account the other fixed and random factors. In order to identify the acoustic similarity between human vocalisations and those of chimpanzees, bonobos and macaques, we performed relevant pairwise comparisons on Mahalanobis distances from the centroid of Human vocalizations: for each affect, we compared: Human vs Chimpanzee, Human vs Bonobo; Human vs Macaque; Chimpanzee vs Bonobo; Chimpanzee vs Macaque and Bonobo vs Macaque. Hence, each subset of data (e.g. threat chimpanzee) appeared a maximum total of 3 times in the pairwise comparisons, leading us to compare our p-values to Bonferroni corrected alpha level of P_corrected_ = .05/3 = .017.

#### Vocal recognition performance

First, we investigated if participants’ recognition accuracy in the categorisation and discrimination tasks was significantly above chance for each affect per species (i.e. three affects x 4 species = 12 separate tests). Per participant, we calculated the proportion of correct answers for each affect-species set of calls (N = 8 calls in each set) and then used one-sample t-tests to examine whether proportion of correct answers was significantly above chance per task (0.33 for categorization task; 0.5 for discrimination task).

Next, to test our hypotheses of phylogenetic distance (hypothesis 1); acoustic similarity (hypothesis 2) or a combination of both (hypothesis 3), we ran GLMMs for both categorization and discrimination tasks separately to examine whether species and affect predicted participant accuracy expressed as the number of correct answers for each type of stimulus (species*affect e.g. chimpanzee distress). We first tested the models against a null model containing only intercept and random effects. All GLMMs were fitted by REML on R.studio using the “bobyqa” function (optimization by quadratic approximation with a set maximum of 1’000’000 iterations) and the link “logit” for a standard logistic distribution of errors and a binomial distribution including: Species (human, chimpanzee, bonobo, and rhesus macaque) and Affect (affiliative, threat, and distress) as fixed factors, accuracy in either the discrimination or categorization task as the Response Variable and participant IDs as random factor.

To relate our results with the acoustic analyses, we ran the same contrasts, i.e. Human vs Chimpanzee, Human vs Bonobo; Chimpanzee vs Bonobo; Chimpanzee vs Macaque and Bonobo vs Macaque for each affect.

## Results

### Acoustic analyses

The DA allowed us to compute Mahalanobis distances for all stimuli compared to Human vocalizations (Figure 2). A GLMM analysis on Mahalanobis distances revealed the full model including main effects and the interaction between Distance-Species and Affect explained significantly more variance compared to the null model (χ^2^(11) = 120.2, p < 0.001).

**Figure 2:**
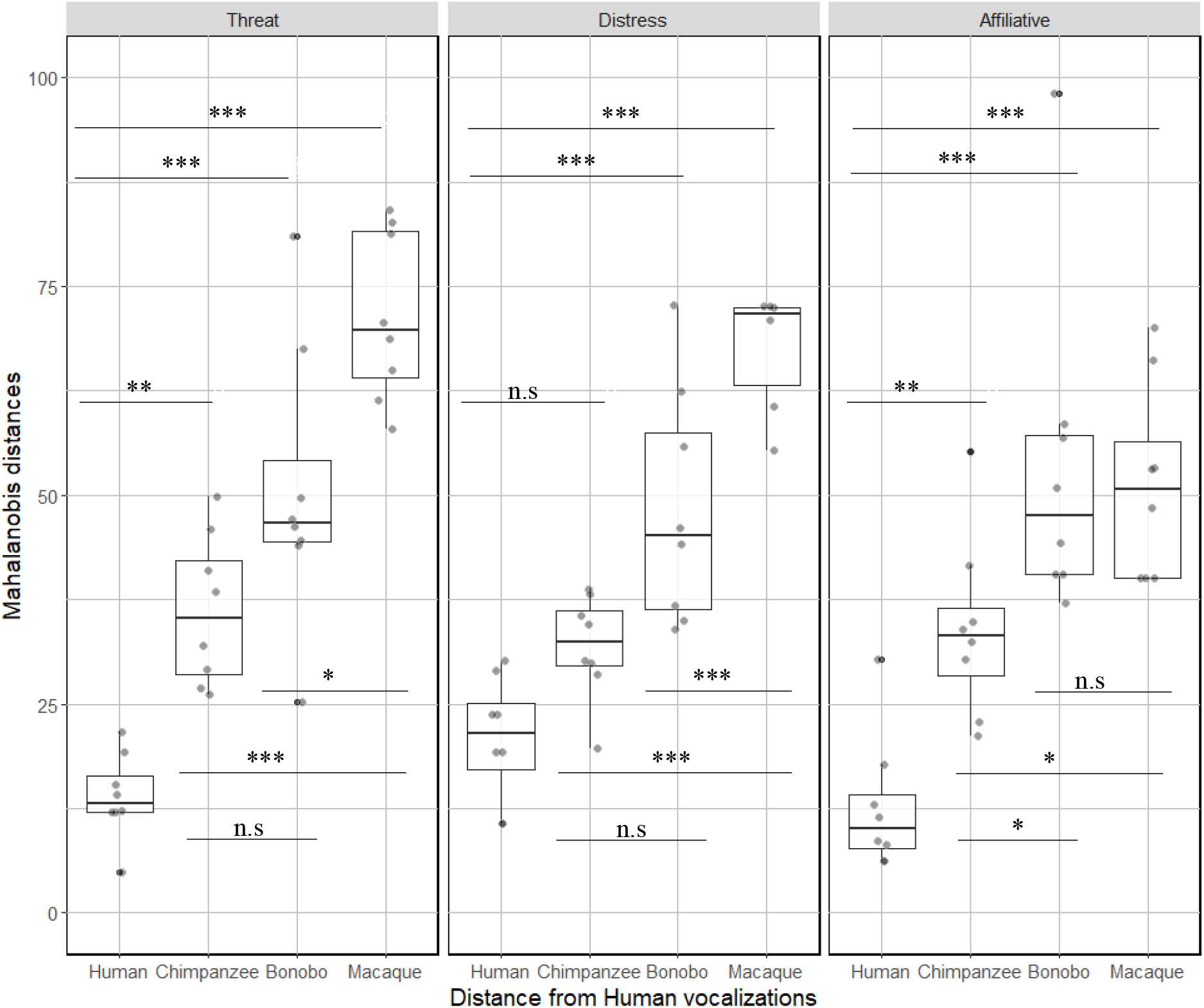
Boxplot of Mahalanobis distances for the 96 vocalizations representing acoustic distances from human voice compared to the other species vocalizations for the different affective states. Higher values represent greater acoustic distances. (*\* <0*.*017; **<0*.*003; ***<0*.*0003)*.

The contrasts in the full model for the comparisons between the levels of distance from the Human Centroid (Distance-Species) for each level of Affects are reported in Table 2 (see also Figure 2). When corrected for multiple comparisons, pairwise comparisons revealed that Mahalanobis distances to human centroids for human vocalisations were significantly smaller than for all bonobo and all macaque vocalisations, as well as affiliative and threat chimpanzee vocalizations, but not chimpanzee distress calls. Chimpanzee and bonobo vocalizations (when plotted from human vocalization centroids) were not significantly different at the levels of distress and threat, but bonobo affiliative vocalisations were significantly further from the human centroid than chimpanzee affiliative vocalisations (see Table 2; Figure 2). Macaque vocalisations were significantly further from the human centroid than chimpanzee vocalisations for all affects. Macaque vocalisations were significantly further from the human centroid than bonobo vocalisations for threat and distress calls, but not affiliative calls.

**Table 1:** Table summarizing the statistical values for the GLMM of acoustic Mahalanobis distances including main effects and the interaction.

| Summary of the model for acoustic Mahalanobis distances | Df | F value | p-value |
| --- | --- | --- | --- |
| Distance-Species | 3 | 62.75 | <.001 |
| Affect | 2 | 2.55 | .084 |
| Distance-Species:Affect | 6 | 2.74 | .018 |

**Table 2:** Table summarizing the results of pairwise comparisons in GLMMs for acoustic across species (Chimpanzee, Bonobo and Macaque) and affect (Threat, distress, affiliative). All p-values are compared to a corrected alpha level of 0.017 (*\* <0*.*017; **<0*.*003; ***<0*.*0003)*. Abbreviations: (Mac) Macaque; (Chimp) Chimpanzee; (affiliat.) affiliative.

|  | *Chimp threat* | *Bonobo threat* | *Mac threat* |  | *Chimp distress* | *Bonobo distress* | *Mac distress* |  | *Chimp affiliat.* | *Bonobo affiliat.* | *Mac affiliat.* |
| --- | --- | --- | --- | --- | --- | --- | --- | --- | --- | --- | --- |
| *Human threat* | $\chi^2(1)=10.3$;<br>p=0.001<br>** | $\chi^2(1)=28.2$;<br>p<0.001<br>*** | $\chi^2(1)=69.23$;<br>p<0.001<br>*** | *Human distress* | $\chi^2(1)=2.58$; p=.11 | $\chi^2(1)=15.8$;<br>p<0.001<br>*** | $\chi^2(1)=75.72$;<br>p<0.001<br>*** | *Human affiliat.* | $\chi^2(1)=9.52$;<br>p=0.002<br>** | $\chi^2(1)=34.5$;<br>p<0.001<br>*** | $\chi^2(1)=31.35$;<br>p<0.001<br>*** |
| *Chimp threat* | -- | $\chi^2(1)=4.42$;<br>p=0.036 | $\chi^2(1)=26.0$;<br>p<0.001<br>*** | *Chimp distress* | | $\chi^2(1)=5.67$;<br>p=0.017 | $\chi^2(1)=50.36$;<br>p<0.001<br>*** | *Chimp affiliat.* | -- | $\chi^2(1)=7.79$;<br>p=0.005<br>* | $\chi^2(1)=6.32$;<br>p=0.012<br>* |
| *Bonobo threat* | | -- | $\chi^2(1)=9.02$;<br>p=0.003<br>* | *Bonobo distress* | | -- | $\chi^2(1)=22.2$;<br>p<0.001<br>*** | *Bonobo affiliat.* | | -- | $\chi^2(1)=0.08$; p=0.78 |

Overall, while the pattern of Mahalanobis distances from the human centroid for threat vocalizations appears to mirror phylogenetic distance between species (with H > C=B > M), we found significant variation for both distress and affiliative vocalizations. With respect to distress calls, the pattern suggests that great ape calls are acoustically similar to each other, but different from macaque calls (H=C=B>M). In contrast, human affiliative calls are significantly different from all other calls, with chimpanzee calls being significantly closer to the human centroid than bonobo or macaque calls (H>C>B=M). The statistical analysis for all other comparisons can be found in the Supplementary Material.

### Vocal recognition performance

Patterns of performance against chance level, as well as between species and affect, differed for categorisation and discrimination.

#### Categorization

Participants were above chance for detecting affect for both human and chimpanzee vocalizations; this was also the case for assigning distress and affiliative calls for bonobos, but not threat calls. In contrast, no call type reached significance for macaques (Figure 3).

**Figure 3:**
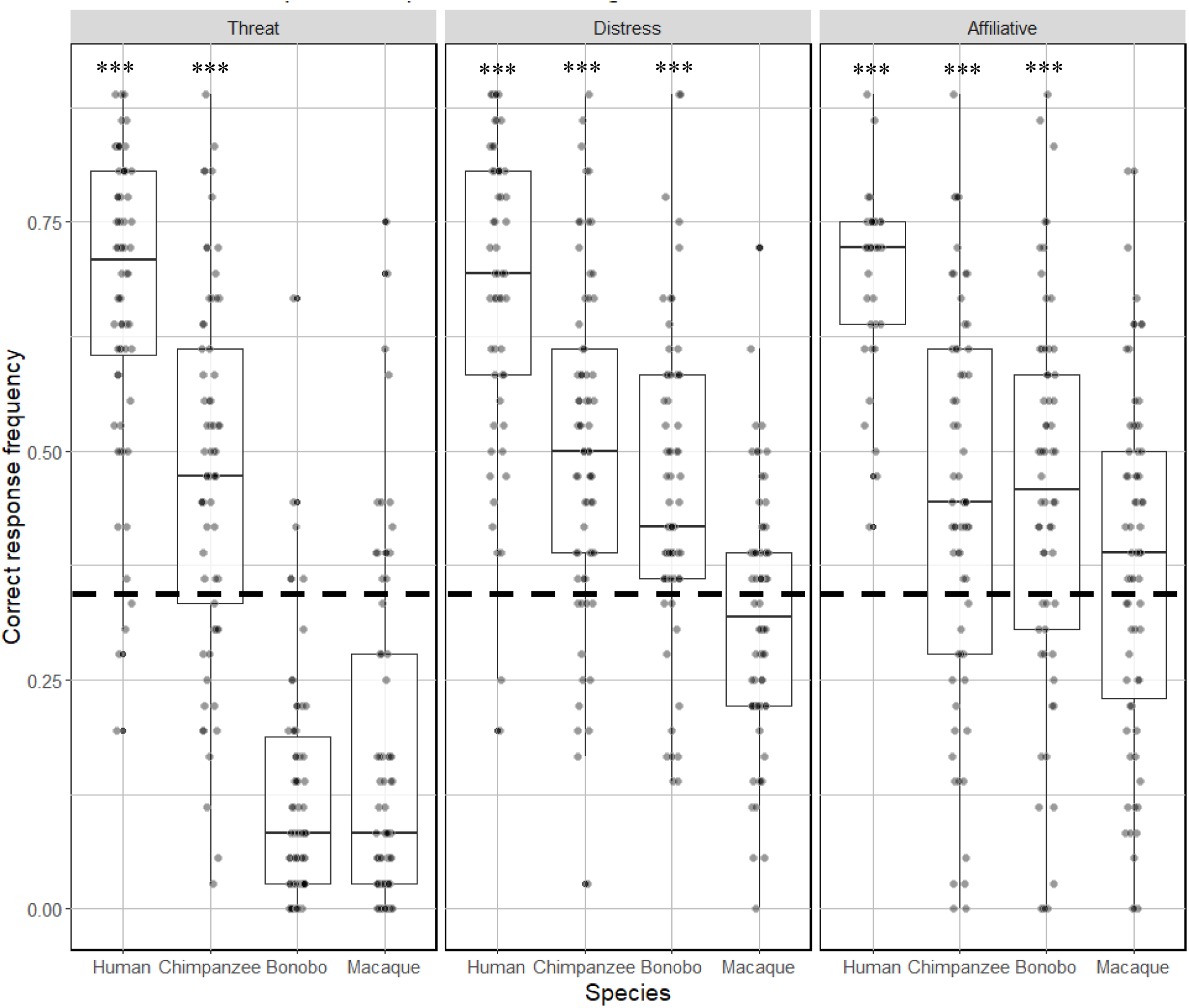
Boxplot illustrating the proportion of correct responses for each category of stimuli in the Categorization task. Higher values represent greater accuracy. One sample T-test analyses against chance level (0.33 - represented with the dotted line) are shown. Note that all types of stimuli were categorized at a level significantly above chance, with the exception of all macaque calls and threatening bonobo calls. See Table S3 in Sup Mat for the summary of the t-values testing whether participants’ accuracy was above chance level. *** p < 0.001.

A GLMM comparison for the categorization task between the null model and the full model with main effects and the interaction (Species and Affects) revealed the full model explained a significant amount of variance in the data χ^2^(11) = 609.3, p < 0.001, see Table 3).

**Table 3:** Table summarizing the main values for GLMMs of accuracy for the Categorization task according to main factors and the interaction.

| Full model for Categorization | Df | Chi-squared | p-value |
| --- | --- | --- | --- |
| Species | 3 | 234.92 | <.001 |
| Affects | 2 | 64.62 | <.001 |
| Species*Affects | 6 | 17.23 | <.001 |

Contrast analysis revealed that human vocalizations were systematically better recognized than chimpanzee, bonobo and macaque vocalizations across all levels of affect (Table 4). In contrast, accuracy with chimpanzee and bonobos distress and affiliative calls was similar, with chimpanzee threat calls being more accurately categorised than bonobo threat calls. Chimpanzee and bonobo distress and affiliative calls were both more accurately categorised than macaque calls. However, macaque threat calls were more accurately categorised than bonobo threat calls. All contrasts are reported in Table 4. Note that all contrasts were compared to a corrected P for multiple comparisons (Bonferroni correction: P_corrected_ = .05/3=.017).

**Table 4:** Table summarizing the results of pairwise comparisons in GLMMs for categorization across species (Chimpanzee, Bonobo and Macaque) and affect (Threat, distress, affiliative. All p-values are compared to a corrected alpha level of 0.017 (*\* <0*.*017; **<0*.*003; ***<0*.*0003)*. Abbreviations: (Mac) Macaque; (Chimp) Chimpanzee; (affiliat.) affiliative.

|  | *Chimp threat* | *Bonobo threat* | *Mac threat* |  | *Chimp distress* | *Bonobo distress* | *Mac distress* |  | *Chimp affiliat.* | *Bonobo affiliat.* | *Mac affiliat.* |
| --- | --- | --- | --- | --- | --- | --- | --- | --- | --- | --- | --- |
| **Human threat** | $\chi^2(1)=37.37$; $p<0.001$ *** | $\chi^2(1)=304.97$; $p<0.001$ *** | $\chi^2(1)=252.77$; $p<0.001$ *** | **Human distress** | $\chi^2(1)=44.49$; $p<0.001$ *** | $\chi^2(1)=56.13$; $p<0.001$ *** | $\chi^2(1)=158.69$; $p<0.001$ *** | **Human affiliat.** | $\chi^2(1)=132.47$; $p<0.001$ *** | $\chi^2(1)=122.59$; $p<0.001$ *** | $\chi^2(1)=200.93$; $p<0.001$ *** |
| **Chimp threat** | -- | $\chi^2(1)=128.84$; $p<0.001$ *** | $\chi^2(1)=95.77$; $p<0.001$ *** | **Chimp distress** | | $\chi^2(1)=0.68$; $p=0.41$ | $\chi^2(1)=35.13$; $p<0.001$ *** | **Chimp affiliat.** | -- | $\chi^2(1)=0.19$; $p=0.66$ | $\chi^2(1)=7.10$; $p<0.008$ * |
| **Bonobo threat** | | -- | $\chi^2(1)=2.45$; $p<0.12$ | **Bonobo distress** | | -- | $\chi^2(1)=26.06$; $p<0.001$ *** | **Bonobo affiliat.** | | -- | $\chi^2(1)=9.63$; $p<0.002$ ** |

#### Discrimination

Participants were above chance when detecting affect for both human and chimpanzee vocalizations; this was also the case for assigning distress and affiliative calls for bonobos and macaque calls. However, threat calls for the two latter species were not discriminated at a level significantly above chance (Figure 4).

**Figure 4:**
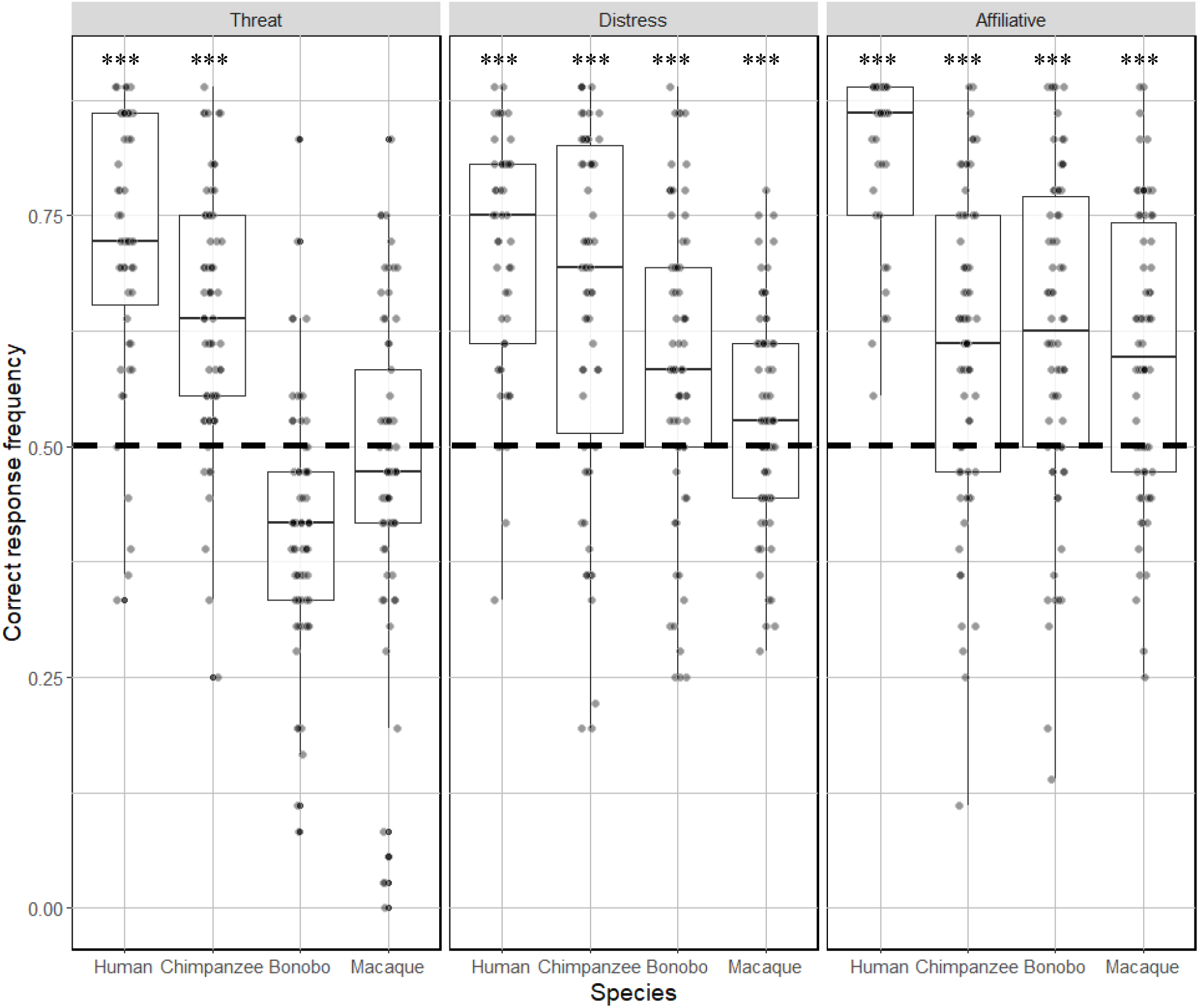
Boxplot illustrating the proportion of correct responses in the Discrimination task. Higher values represent greater accuracy. One sample T-test analyses against chance level (0.5 - shown with the dotted line) are reported. Note that all types of stimuli were discriminated at above chance levels with the exception of all macaque calls and threatening bonobo calls. *** p < 0.001.

A GLMM run on the discrimination task data revealed that the full model explained significantly more variation in the data than the null model χ^2^(11) = 436.97, p < 0.001, see Table 5).

**Table 5:** Table summarizing the main values for GLMMs of accuracy for the Discrimination task according to main factors and the interaction.

| Full model for Discrimination | Df | Chi-squared | p-value |
| --- | --- | --- | --- |
| Species | 3 | 150.62 | <.001 |
| Affect | 2 | 32.52 | <.001 |
| Species*Affect | 6 | 12.23 | <.001 |

Contrast analysis revealed that human vocalizations were systematically better recognized than chimpanzee, bonobo and macaque vocalizations at all levels of affect (Table 6). Chimpanzee threat calls were significantly better discriminated compared to threat calls of both bonobo and macaques, whilst macaque threat calls were better discriminated than bonobo calls. In contrast, while participants were again significantly better at discriminating chimpanzee distress vocalizations than bonobo and macaque distress vocalizations, bonobo distress calls were discriminated better than macaque vocalizations. Finally, none of the contrasts reached significance level for comparison of affiliative vocalizations in non-human primates.

**Table 6:**
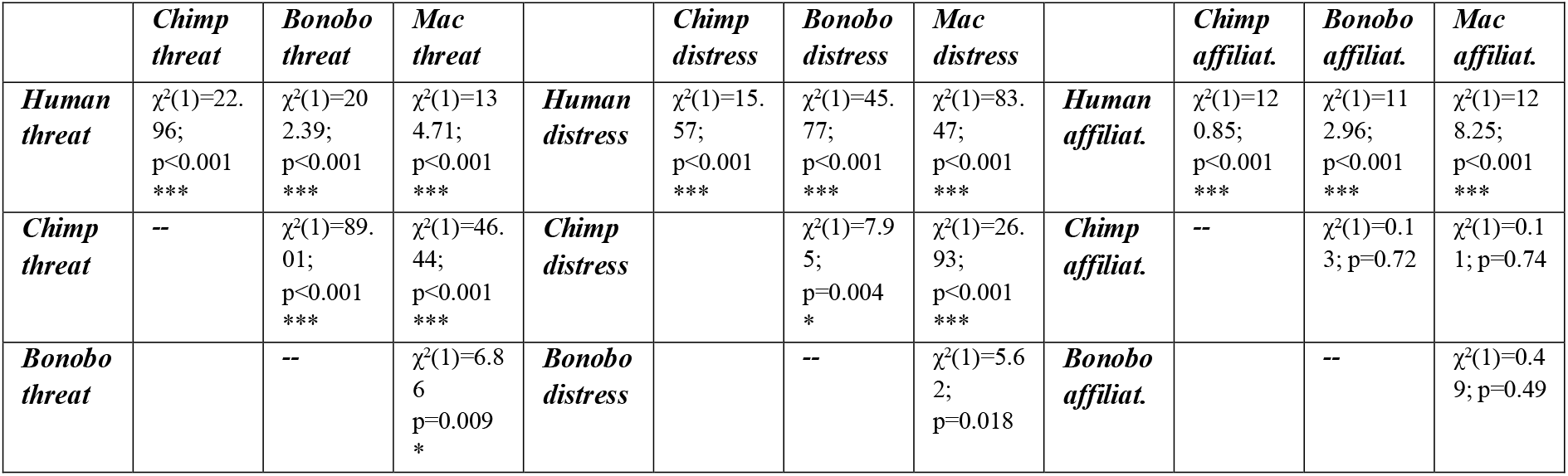
Table summarizing the results of pairwise comparisons in GLMMs for discrimination across species (Chimpanzee, Bonobo and Macaque) and affect (Threat, distress, affiliative. All p-values are compared to a corrected alpha level of 0.017 (*\* <0*.*017; **<0*.*003; ***<0*.*0003)*. Abbreviations: (Mac) Macaque; (Chimp) Chimpanzee; (affiliat.) affiliative.

## Discussion

In this study, we used a combination of acoustic analyses and experimental recognition tasks to investigate how humans perceive primate vocal communication of affect. Using acoustic analysis, we examined the extent to which phylogenetic proximity and the category of affect (threat, distress, affiliative) predicted call acoustic similarity in human, chimpanzee, bonobo and rhesus macaque calls. Using these acoustic analyses, we then tested whether phylogenetic similarity (hypothesis 1), acoustic distance (hypothesis 2) or a combination of both (hypothesis 3) best explained human recognition of affect in these primate vocalisations. Results from two subsequent recognition tasks - discrimination and categorization - which varied on task difficulty, demonstrated that participants were generally better at categorizing and discriminating human and chimpanzee vocalizations versus bonobo and rhesus macaque calls, supporting our third hypothesis both that phylogenetic distance and acoustic similarity might influence human recognition accuracy. There was however more variation for bonobo calls, with participants having difficulty recognizing their threat calls. Finally, macaque calls were the least recognized of all primate vocalizations tested, consistent with a phylogenetic distance hypothesis.

In terms of the acoustic analyses, the acoustic factors extracted in our Discriminant Analyses revealed the crucial role of specific acoustic features such as spectral, frequency, and loudness parameters (see Supp Mat) to distinguish affective vocalizations expressed by different primate species. Our analysis of Mahalanobis distances showed that overall, human vocalizations in the three selected affect categories were acoustically closest to chimpanzee vocalizations, with distress calls virtually indistinguishable by our model. By contrast, overlap with bonobo calls was much lower, despite chimpanzees and bonobos being equally phylogenetically related to humans. Affiliative bonobo vocalizations also showed significant differences in acoustic structure from those of chimpanzees but not from those of macaques, despite chimpanzees being much more closely related to them. Note however that macaque calls were also not significantly acoustically different from chimpanzee affiliative calls. The variation outlined between chimpanzee and bonobo calls is in line with current evidence that despite their genetic proximity, the two species have known behavioural (Gruber & Clay, 2016), neurological (Staes et al., 2018) and morphological differences, including a shorter larynx for bonobos, which drives a higher F0 in their vocalizations (Grawunder et al., 2018). Overall, the phylogenetic hypothesis (H<C=B<M) was only partially supported by the distance pattern found for threat vocalizations, while the rest of the affective contexts offered a mixed bag of patterns, distress grouping apes together (including humans), and affiliative mostly singling out human calls.

Importantly, the acoustic similarity of chimpanzee, bonobo and rhesus monkey vocalizations to those of humans did not reliably predict participants’ ability to categorize and discriminate their affective content. Although more accurate categorization of human vocal affect was to be expected, participants were nonetheless better than chance for detecting the affective content of most vocalizations of each ape species, apart from bonobo threat calls. Crucially, the latter calls had been characterized as similar by the Mahalanobis analysis, suggesting that additional factors come in play when recognizing primate calls. Similarly, despite the lack of acoustic differences between macaque affiliative calls and other great ape affiliative vocalizations, participants struggled to accurately categorize and discriminate their affective content. A possibility to explain these results is that we do not know which of the acoustic factors measured are the most attended to by humans; possibly skewing the weight that can be given to each parameters and making their application to vocalizations that differ substantially from human calls harder; further work will therefore have to fine-tune an acoustic toolbox designed for human vocalisations to phylogenetically close species calls that nonetheless differ acoustically from our own vocalizations. Yet, these findings should not overcast the fact that our participants were generally good at classifying primate calls, particularly ape calls, with the exception of bonobo threat calls. Finally, the results for rhesus macaque calls underline task differences with participants above chance level for discriminating between affiliative and distress calls, with the former being closest to apes’ vocalizations in the Mahalanobis analysis, but not the latter. This underlines once again that while acoustic distance may help participants to correctly classify calls in some contexts, there may be no relation in other contexts, suggesting the existence of additional factors.

Results from this study complement previous research showing highly mixed performance for detecting the affective nature of rhesus monkey calls (Fritz et al., 2018; Scheumann et al., 2014; Scheumann, Hasting, Zimmermann, & Kotz, 2017; Belin, Fecteau, et al., 2008) (Linnankoski et al., 1994). Interestingly, our study also outlines that the differences in findings may be due to the task required from the participants. Both our study and that of Linnankoski and colleagues’, which found some recognition of macaque affective calls, used a forced-choice method (the use of two or more specific response options) to identify affective cues, whereas other studies used Likert response scales. Overall, discrimination led to a higher recognition for participants compared to categorization, with participants only failing to recognize threat calls in bonobos and macaques. This may be due to the fact that categorization is itself more complicated cognitively than discrimination (with three options rather than two), a phenomenon already described when solely using human emotional calls (Dricu et al. 2017; Gruber et al. 2020). Conversely, the difference between the performances in the tasks also means that categorization tasks may be more discriminatory in pointing out the factors that affect most the identification of the correct affect. Compared to discrimination, where patterns of responses do not underline a particular hypothesis, we found that patterns of categorization in the GLMM for distress and affective calls followed a phylogenetic pattern (H>C=B>M), while the overall frequency in performance suggested an acoustic pattern for distress only (H=C=B>M). The result for distress in particular highlights that both acoustic and phylogenetic factors can be identified separately for the same affect, showing the complexity of the recognition process overall; but also that categorization tasks rather than discrimination tasks or Likert scales may offer the granularity necessary to identify the different intervening factors.

## Conclusion

Overall, we demonstrated the ability of humans to both categorize and discriminate affective cues in other primate species’ vocalizations, although we found contextual differences across species and affect, which are not readily explained either by phylogeny or acoustic differences. Beyond single explanations, by using the acoustic distance between four primate species with varying levels of phylogenetic similarity whose vocalisations also varied in different ways with respect to acoustic similarity across affect categories, our study demonstrates that the perception of emotional cues by humans in primate vocalizations is a complex process that does not solely rely on phylogenetic or acoustic similarity. In particular, the inclusion of bonobo vocalizations, while not allowing us to disentangle phylogeny from acoustic factors, underlines the idiosyncratic evolutionary pathway on which they have engaged compared to chimpanzees (Grawunder et al., 2018), and also suggests that there are acoustic factors partially independent from phylogeny and affective content that influence the recognition of calls in NHPs. In this light, bonobo calls were most often verbally pointed out by participants as the most unusual. Therefore, the unfamiliarity of naïve participants with some vocalizations (e.g. bonobo threatening calls) could be at play. Hence, future work will need to additionally disentangle the effect of familiarity from potential acoustic parameters. It would also be interesting to explore neural correlates associated with these phylogenetic and acoustic parameters, to offer another level analysis to the behavioural differences outlined in the present study. Finally, we hope that these new findings will contribute to a better understanding of emotional processing origin in humans, by highlighting where the treatment of both primate and human emotions is similar, and where our own species has differed during its evolution.

## Acknowledgements

We thank Katie Slocombe very much for providing some of the auditory stimuli as well as extensive comments on former versions of this preprint. We thank the Swiss National Science foundation (SNSF) for supporting this interdisciplinary project (CR13I1_162720 / 1 – DG-TG), and the Swiss Center for Affective Sciences. ZC has received support from the ESRC-ORA (ES/S015612/1), the ERC Starting Grant (802979, and CD from the foundation Ernst and Lucie Schmidheiny. TG was additionally supported by a grant of the SNSF during the final re-writing of this article (grant PCEFP1_186832).

## References

Anderson, D. J., & Adolphs, R. (2014). A Framework for Studying Emotions Across Phylogeny. Cell, 157(1), 187–200. https://doi.org/10.1016/j.cell.2014.03.003

Arnal, L. H., Flinker, A., Kleinschmidt, A., Giraud, A.-L., & Poeppel, D. (2015). Human Screams Occupy a Privileged Niche in the Communication Soundscape. Current Biology : CB, 25(15), 2051–2056. https://doi.org/10.1016/j.cub.2015.06.043

Bates, D., Mächler, M., Bolker, B., & Walker, S. (2015). Fitting Linear Mixed-Effects Models Using lme4. Journal of Statistical Software, 67(1), 1–48. https://doi.org/10.18637/jss.v067.i01

Belin, P., Fecteau, S., Charest, I., Nicastro, N., Hauser, M. D., & Armony, J. L. (2008). Human cerebral response to animal affective vocalizations. Proceedings. Biological Sciences, 275(1634), 473–481. https://doi.org/10.1098/rspb.2007.1460

Belin, P., Fillion-Bilodeau, S., & Gosselin, F. (2008). The Montreal Affective Voices: A validated set of nonverbal affect bursts for research on auditory affective processing. Behavior Research Methods, 40(2), 531–539. https://doi.org/10.3758/BRM.40.2.531

Briefer, E. (2012). Vocal Expression of Emotions in Mammals: Mechanisms of Production and Evidence. Communication Skills. https://animalstudiesrepository.org/comski/1

Brunswick, E. (1956). Perception and the representative design of psychological experiments. University of California Press.

Davila Ross, M., Owren, M. J., & Zimmermann, E. (2009). Reconstructing the evolution of laughter in great apes and humans. Current Biology: CB, 19(13), 1106–1111. https://doi.org/10.1016/j.cub.2009.05.028

Dricu, M., Ceravolo, L., Grandjean, D., & Frühholz, S. (2017). Biased and unbiased perceptual decision-making on vocal emotions. Scientific Reports, 7(1), 16274. https://doi.org/10.1038/s41598-017-16594-w

Eyben, F., Scherer, K., Schuller, B., Sundberg, J., André, E., Busso, C., Devillers, L., Epps, J., Laukka, P., Narayanan, S., & Truong, K. P. (2016). The Geneva Minimalistic Acoustic Parameter Set (GeMAPS) for Voice Research and Affective Computing. IEEE transactions on affective computing, 7(2), 190–202. https://doi.org/10.1109/TAFFC.2015.2457417

Ferdenzi, C., Patel, S., Mehu-Blantar, I., Khidasheli, M., Sander, D., & Delplanque, S. (2013). Voice attractiveness: Influence of stimulus duration and type. Behavior Research Methods, 45(2), 405–413. https://doi.org/10.3758/s13428-012-0275-0

Ferry, A. L., Hespos, S. J., & Waxman, S. R. (2013). Nonhuman primate vocalizations support categorization in very young human infants. Proceedings of the National Academy of Sciences, 110(38), 15231–15235. https://doi.org/10.1073/pnas.1221166110

Filippi, P. (2016). Emotional and Interactional Prosody across Animal Communication Systems: A Comparative Approach to the Emergence of Language. Frontiers in Psychology, 7, 1393. https://doi.org/10.3389/fpsyg.2016.01393

Filippi, P., Congdon, J. V., Hoang, J., Bowling, D. L., Reber, S. A., Pašukonis, A., Hoeschele, M., Ocklenburg, S., de Boer, B., Sturdy, C. B., Newen, A., & Güntürkün, O. (2017). Humans recognize emotional arousal in vocalizations across all classes of terrestrial vertebrates: Evidence for acoustic universals. Proceedings of the Royal Society B: Biological Sciences, 284(1859), 20170990. https://doi.org/10.1098/rspb.2017.0990

Frijda, N. H. (1987). Emotion, cognitive structure, and action tendency. Cognition and Emotion, 1(2), 115–143. https://doi.org/10.1080/02699938708408043

Frijda, N. H. (2016). The evolutionary emergence of what we call “emotions.” Cognition and Emotion, 30(4), 609–620. https://doi.org/10.1080/02699931.2016.1145106

Fritz, T., Mueller, K., Guha, A., Gouws, A., Levita, L., Andrews, T. J., & Slocombe, K. E. (2018). Human behavioural discrimination of human, chimpanzee and macaque affective vocalisations is reflected by the neural response in the superior temporal sulcus. Neuropsychologia, 111, 145–150. https://doi.org/10.1016/j.neuropsychologia.2018.01.026

Grandjean, D., Bänziger, T., & Scherer, K. R. (2006). Intonation as an interface between language and affect. In Progress in Brain Research (Vol. 156, pp. 235–247). Elsevier. https://doi.org/10.1016/S0079-6123(06)56012-1

Grandjean, D., Sander, D., Pourtois, G., Schwartz, S., Seghier, M. L., Scherer, K. R., & Vuilleumier, P. (2005). The voices of wrath: Brain responses to angry prosody in meaningless speech. Nature Neuroscience, 8(2), 145–146. https://doi.org/10.1038/nn1392

Grawunder, S., Crockford, C., Clay, Z., Kalan, A. K., Stevens, J. M. G., Stoessel, A., & Hohmann, G. (2018). Higher fundamental frequency in bonobos is explained by larynx morphology. Current Biology: CB, 28(20), R1188–R1189. https://doi.org/10.1016/j.cub.2018.09.030

Gross, J. J. (1998). The emerging field of emotion regulation: An integrative review. Review of General Psychology, 271–299.

Gruber, T., & Clay, Z. (2016). A Comparison Between Bonobos and Chimpanzees: A Review and Update. Evolutionary Anthropology: Issues, News, and Reviews, 25(5), 239–252. https://doi.org/10.1002/evan.21501

Gruber, T., & Grandjean, D. M. (2017). A comparative neurological approach to emotional expressions in primate vocalizations. Neuroscience and Biobehavioral Reviews, 73, 182–190.

Kamiloğlu, R. G., Slocombe, K. E., Haun, D. B. M., & Sauter, D. A. (2020). Human listeners’ perception of behavioural context and core affect dimensions in chimpanzee vocalizations. Proceedings of the Royal Society B: Biological Sciences, 287(1929), 20201148. https://doi.org/10.1098/rspb.2020.1148

Kelly, T., Reby, D., Levréro, F., Keenan, S., Gustafsson, E., Koutseff, A., & Mathevon, N. (2017). Adult human perception of distress in the cries of bonobo, chimpanzee, and human infants. Biological Journal of the Linnean Society, 120(4), 919–930. https://doi.org/10.1093/biolinnean/blw016

Kret, M. E., Prochazkova, E., Sterck, E. H. M., & Clay, Z. (2020). Emotional expressions in human and non-human great apes. Neuroscience & Biobehavioral Reviews. https://doi.org/10.1016/j.neubiorev.2020.01.027

Linnankoski, I., Laakso, M., Aulanko, R., & Leinonen, L. (1994). Recognition of emotions in macaque vocalizations by children and adults. Language & Communication, 14(2), 183–192. https://doi.org/10.1016/0271-5309(94)90012-4

Mahalanobis, P. C. (1936). On the Generalized Distance in statistics. In Proceedings of National Institute of Sciences (Vol. 2, p. 49.55).

Manser, M. B. (2001). The acoustic structure of suricates’ alarm calls varies with predator type and the level of response urgency. Proceedings of the Royal Society B: Biological Sciences, 268(1483), 2315–2324. https://doi.org/10.1098/rspb.2001.1773

Morton, E. S. (1977). On the Occurrence and Significance of Motivation-Structural Rules in Some Bird and Mammal Sounds. The American Naturalist, 111(981), 855–869. https://doi.org/10.1086/283219

Morton, E. S. (1982). Grading, discreteness, redundancy, and motivation-structural rules. In Acoustic Communication in Birds (Kroodsma, D.E., Miller, E.H. and Ouellet, H., pp. 182–212). Academic Press.

Nesse, R. M. (1990). Evolutionary explanations of emotions. Human Nature, 1(3), 261–289. https://doi.org/10.1007/BF02733986

Sander, D., Grandjean, D., Pourtois, G., Schwartz, S., Seghier, M. L., Scherer, K. R., & Vuilleumier, P. (2005). Emotion and attention interactions in social cognition: Brain regions involved in processing anger prosody. NeuroImage, 28(4), 848–858. https://doi.org/10.1016/j.neuroimage.2005.06.023

Scherer, K. (2003). Vocal communication of emotion: A review of research paradigms. Speech Communication, 40(1–2), 227–256. https://doi.org/10.1016/S0167-6393(02)00084-5

Scheumann, M., Hasting, A. S., Kotz, S. A., & Zimmermann, E. (2014). The voice of emotion across species: How do human listeners recognize animals’ affective states? PloS One, 9(3), e91192. https://doi.org/10.1371/journal.pone.0091192

Scheumann, M., Hasting, A. S., Zimmermann, E., & Kotz, S. A. (2017). Human Novelty Response to Emotional Animal Vocalizations: Effects of Phylogeny and Familiarity. Frontiers in Behavioral Neuroscience, 11. https://doi.org/10.3389/fnbeh.2017.00204

Schore, J. R., & Schore, A. N. (2008). Modern Attachment Theory: The Central Role of Affect Regulation in Development and Treatment. Clinical Social Work Journal, 36(1), 9–20. https://doi.org/10.1007/s10615-007-0111-7

Staes, N., Smaers, J. B., Kunkle, A. E., Hopkins, W. D., Bradley, B. J., & Sherwood, C. C. (2018). Evolutionary divergence of neuroanatomical organization and related genes in chimpanzees and bonobos. Cortex. https://doi.org/10.1016/j.cortex.2018.09.016

Team, R. (2020). RStudio: Integrated Development for R. RStudio. RStudio, Inc. https://rstudio.com/

Tuttle, R. H. (1993).

Kano, T. 1992. The Last Ape: Pygmy Chimpanzee Behavior and Ecology. Stanford University Press, Stanford, CA, xxviii + 248 pp. ISBN 0-8047-1612-9. Price (hardbound), $45.00. Journal of Mammalogy, 74(1), 239–240. https://doi.org/10.2307/1381928

